# Sources of non-uniform coverage in short-read RNA-Seq data

**DOI:** 10.1101/2025.01.30.634337

**Authors:** Thomas G Brooks, Nicholas F Lahens, Antonijo Mrčela, Jianing Yang, Souparna Purohit, Amruta Naik, Emanuela Ricciotti, Shaon Sengupta, Peter S Choi, Gregory R Grant

## Abstract

The origin of several normal cellular functions and related abnormalities can be traced back to RNA splicing. As such, RNA splicing is currently the focus of a vast array of studies. To quantify the transcriptome, short-read RNA-Seq remains the standard assay. The primary technical artifact of RNA-Seq library prep, which severely interferes with analysis, is extreme non-uniformity in coverage across transcripts. This non-uniformity is present in both bulk and single-cell RNA-Seq and is observed even when the sample contains only full-length transcripts. This issue dramatically affects the accuracy of isoform-level quantification of multi-isoform genes. Understanding the sources of this non-uniformity is critical to developing improved protocols and analysis methods. Here, we explore eight potential sources of non-uniformity. We demonstrate that it cannot be explained by one factor alone. We performed targeted experiments to investigate the effect of fragment length, PCR ramp rate, and ribosomal depletion. We assessed existing data sets with varying sample quality, PCR cycle number, reverse transcriptase, and technical or biological replicates. We found evidence that interference of reverse transcription by secondary structure is unlikely to be the major contributing factor, that rRNA pull-down methods do not cause non-uniformity, that PCR ramp rate does not substantially impact non-uniformity, and that shorter fragments do not reduce non-uniformity. All these findings contradict prior publications or recommendations.

## Introduction

Bulk short read RNA-Seq is a mainstay of modern biology. In a PubMed survey of the 100 most recent publications returned by the search “mouse gene expression,” 49% employ RNA-Seq of some form. Of those, 75% used short read bulk, 28% used single cell (a few used both), none used long reads. At this point, short-read bulk RNA-Seq appears to be a standard tool for transcriptomics indefinitely. Nonetheless, transcript level analysis with short-read bulk RNA-Seq leaves significant room for improvement because of a ubiquitous and severe technical artifact. Figure 1a shows a typical coverage plot of a typical gene (Gapdh). These data come from a high-quality sample where we expect mostly full-length transcripts, yet we observe extreme variability in coverage in the form of wide peaks and valleys. However, in genes with multiple isoforms, these patterns overlap and have been shown to impair isoform-level quantification^2, 3^.

**Figure 1.**
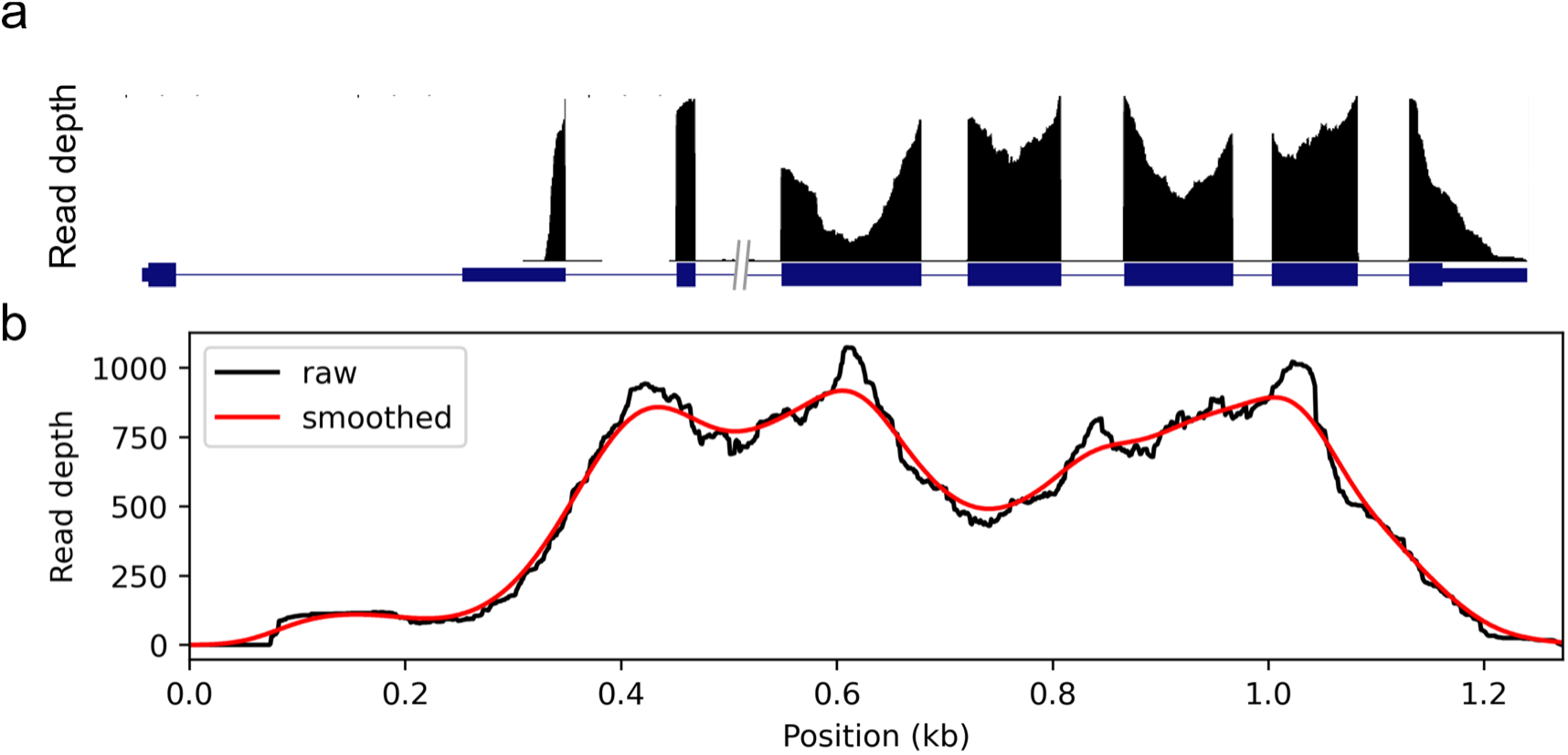
Coverage non-uniformity. (a) A typical RNA-Seq coverage plot showing the peak and valley nature of the signal in the Gapdh gene. (b) The same data shown with as fragment coverage of the transcript. Here, introns are removed, and reads are assumed to be contiguous from one paired end to the other. In red, the same data smoothed (30bp standard deviation Gaussian kernel) to remove small variations.

If this problem can be mitigated, it will enable transcript-level analyses to a level of accuracy that is currently not possible with short-read data. These issues are easily overlooked when working with spreadsheets of quantified data without examining the coverage plots.

The most common library preparation protocols start with ribosomal RNA (rRNA) depletion, followed by fragmentation, reverse transcription, fragment size selection, adapter ligation, and PCR amplification – followed by sequencing. Multiple protocols exist for each of these steps. For example, the typical approach for rRNA depletion is polyA selection, which is very efficient but introduces a significant 3’ bias^4^ and misses any non-polyadenylated RNA. A standard alternative is to hybridize probes to ribosomal sequences and then to remove rRNA by pull-down or digestion. These probe-based approaches avoid the 3’ bias, enabling the sequencing of degraded or fragmented RNA, as well as allowing for sequencing non-polyadenylated RNA, but may introduce their own biases.

Some amount of variability in coverage is due to the inherent variability involved in sequencing a randomly chosen population of fragments. We refer to this as *fragment sampling variability*, however we will show that it is insignificant in anything but low-expressed genes.

There has been considerable effort to algorithmically model and correct for non-uniformity to improve transcript-level quantification, differential expression analysis, and other pursuits. These include alpine^5^, a beta-binomial approach^6^, PennSeq^7^, mseq^8^, and a neural network approach^9^; as well as bias correction models in highly popular quantifiers including Kallisto^10^, Salmon^11^, Cufflinks^12^, and RSEM^13^. These attempt to correct for local sequence biases (such as from hexamer priming), GC content biases, or positional bias (such as 3’ bias from polyA-selected datasets). We have previously found that the bias corrections in Salmon improve quantifications but do not completely correct for non-uniformity in simulated data^14^, with real-world data likely proving even more of a challenge.

Possible sources for bias in RNA-Seq have been reviewed before^15^, and we consulted with Illumina and many researchers and core facilities to identify the most likely factors impacting non-uniform coverage. The explanations given can be divided into eight categories of putative factors: (1) the input RNA sample; (2) fragment sampling variability; (3) random hexamer priming^16^; (4) ribosomal depletion^17^; (5) fragmentation; (6) reverse transcription^18^; (7) PCR^19^; and (8) alignment.

Here, we investigate these putative sources of coverage non-uniformity in turn. We present evidence from public data as well as in-house experiments. We demonstrate that no single factor explains everything, and that all current proposals to improve uniformity fail to fix the problem substantially. We summarize the current progress on this issue; however, the problem is still far from being solved.

## Results

We use data from eight data sets, see Table 1 for descriptions and brief names they will be referred by. Since coverage may contain non-uniformity arising from both technical artifacts as well as from the expression of multiple isoforms, we chose for each data set the 100 highest expressed single-isoform genes where coverage is expected to be uniform. We perform our analyses and plot figures based on these single-isoform genes. Except when intentionally showing introns, we show coverage by concatenating the exons and compute coverage according to fragments so that paired-end reads are connected from both ends even if no bases were read between them, see Figure 1 b. This better captures the fragment library sent to sequencing.

**Table 1.** Data sets Description of the data sets used in this paper.

| **Data set name** | **Organism** | **Tissue** | **Description** | **Accession** | **UMI tags** |
| --- | --- | --- | --- | --- | --- |
| **SEQC** | Human | Universal Human Reference | SEQC samples of UHR (sample A) prepared at four sites | PRJNA208369 | No |
| **GEO** | Mouse | Liver | 6 RiboZero Gold studies from GEO, 2 biological replicate samples from each | GSE77221<br>GSE117134<br>GSE208768<br>PRJNA816471<br>PRJNA788430<br>PRJNA753198 | No |
| **SMART** | Mouse | Hippocampus | Bulk and single-cell SMART-seq data of brain myeloid cells | SRP170969<br>SRP170960 | No |
| **NS1** | Mouse | Liver and heart | Novel experiment with NoSelect samples as well as varying fragment size, and PCR ramp rate | GSE288460 | Yes |
| **NS2** | Mouse | Lung | Novel experiment with biological replicates of NoSelect, polyA, and rRNA pull-down | GSE288461 | Yes |
| **DGRD** | Human | Universal Human Reference | UHR samples that underwent chemical degradation | SRP073267 | No |
| **PCR** | Mouse | Lung | Libraries prepared with varying PCR cycle counts | PRJNA416930 | Yes |
| **TGIRT** | Human | Universal Human Reference | TGIRT-seq libraries, with reverse transcription done at 60°C | PRJNA505326 | No |

Visually, two primary magnitudes of “wiggles” in the coverage plot are evident, see Figure 1 b. It is primarily the larger scale variations, which we refer to as “hills” and “valleys,” which complicate downstream analysis such as isoform quantification.

**Figure 2.**
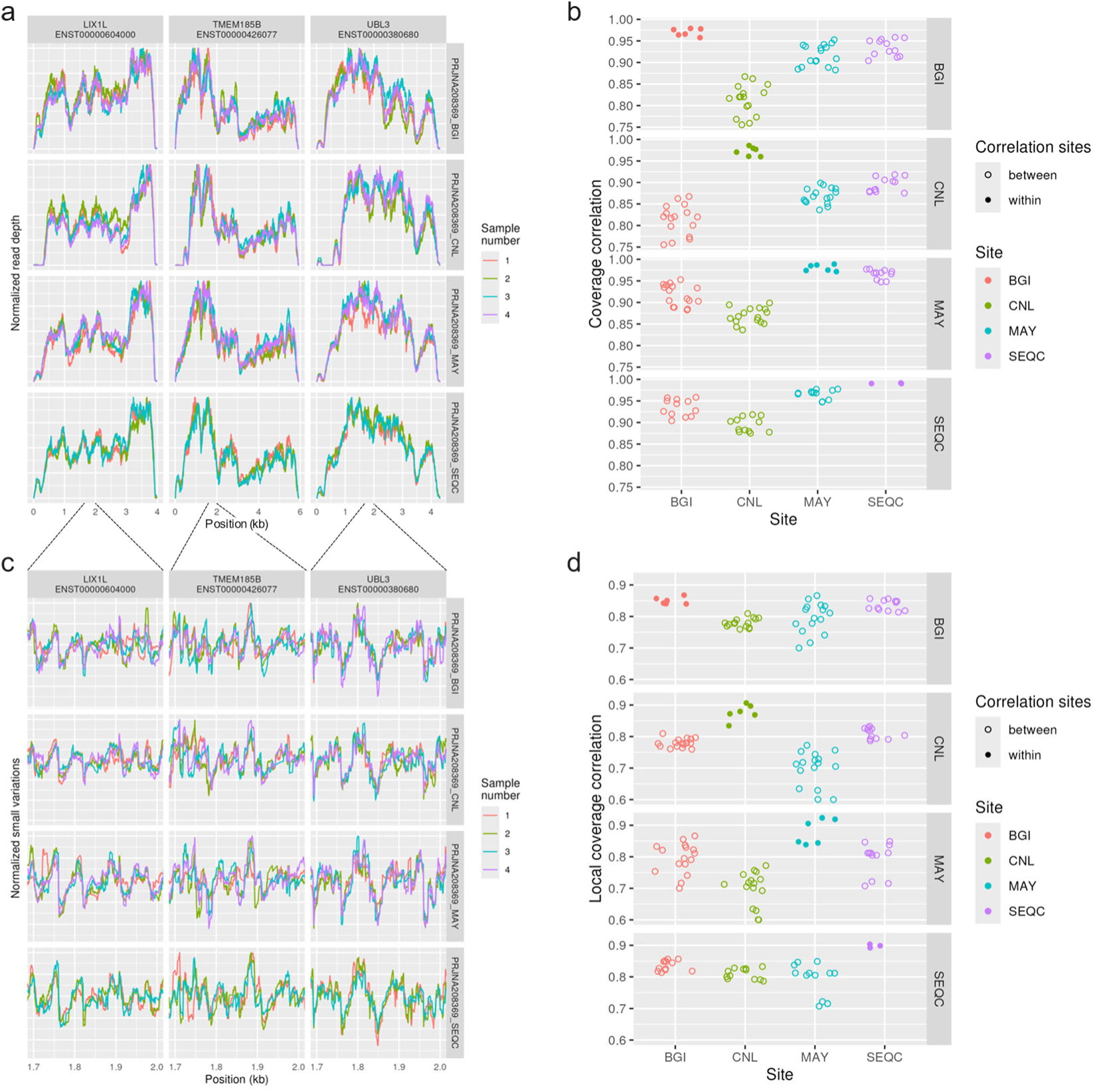
Consistency of non-uniformity across technical replicates. In the SEQC data set, three sites (labelled BGI, CNL, MAY) prepared and sequenced 4 libraries, which were technical replicates of the universal human reference samples. In addition, one library was prepared at a fourth site (labelled SEQC) and then aliquots were sequenced at the BGI, CNL, and MAY sites alongside the libraries prepared at those sites. (a) Coverage in three high-expressed, single-isoform genes is shown. (b) Correlation of coverage across 100 high-expressed, single-isoform genes is shown, after normalizing for read depths, for each pair of samples. (c) Small variations in coverage shown in a 300bp region, computed by removing a smoothed coverage (Gaussian kernel with 30 bp standard deviation). (d) Local correlations, defined as the Pearson correlation of the small variations in coverage after removing smoothed coverage and normalizing for read depths.

To examine technical variation, we used data from SEQC^20^, which includes RNA libraries prepared and sequenced in different batches and at different sites. Technical replicate libraries prepared in the same batch show remarkable consistency in coverage, and sequencing site had little effect, see Figure 2 a-b. Small-scale variations in coverage (after removing smoothed coverage) were highly replicable in technical replicates as well, see Figure 2 c-d.

To examine biological variation, we found six comparable mouse liver studies from the Short Read Archive and chose two replicates from each. Biological replicates within studies show largely similar shapes of their coverage plots, with many showing almost identical traces, see Figure 3 a-b. Across studies non-uniformity was noticeably less consistent. However, certain large drops or peaks appeared to be conserved across most studies. Small-scale variations were again highly replicable within a study, see Figure 3 c-d.

**Figure 3.**
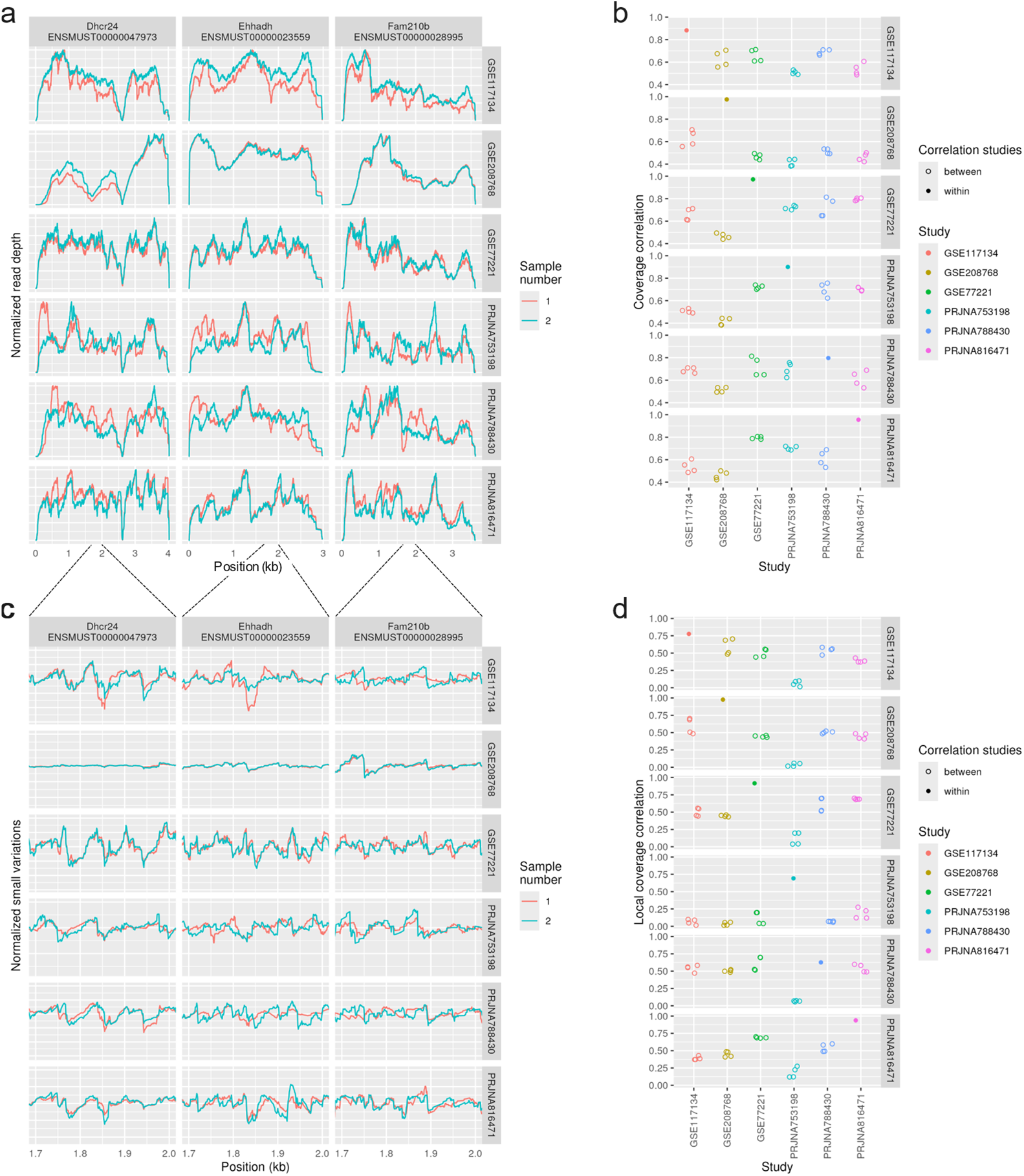
Consistency of non-uniformity across biological replicates and studies. For the GEO data set, we selected two replicates from each of six mouse liver studies available in the Short Read Archive that were prepared with a Ribo-Zero Gold kit (rRNA pull-down) and sequenced on an Illumina sequencer. Normalized read depth coverage of three chosen high-expressed genes with just one annotated isoform is shown. (a) Coverage plots for three select, high-expressed single-isoform genes. (b) Correlation of coverage across 100 high-expressed, single-isoform genes is shown, after normalizing for read depths, for each pair of samples. (c) Small variations in coverage shown in a 300bp region, computed by removing a smoothed coverage (Gaussian kernel with 30 bp standard deviation). (d) Local correlations, defined as the Pearson correlation of the small variations in coverage after removing smoothed coverage, after normalizing for read depths.

**Figure 4.**
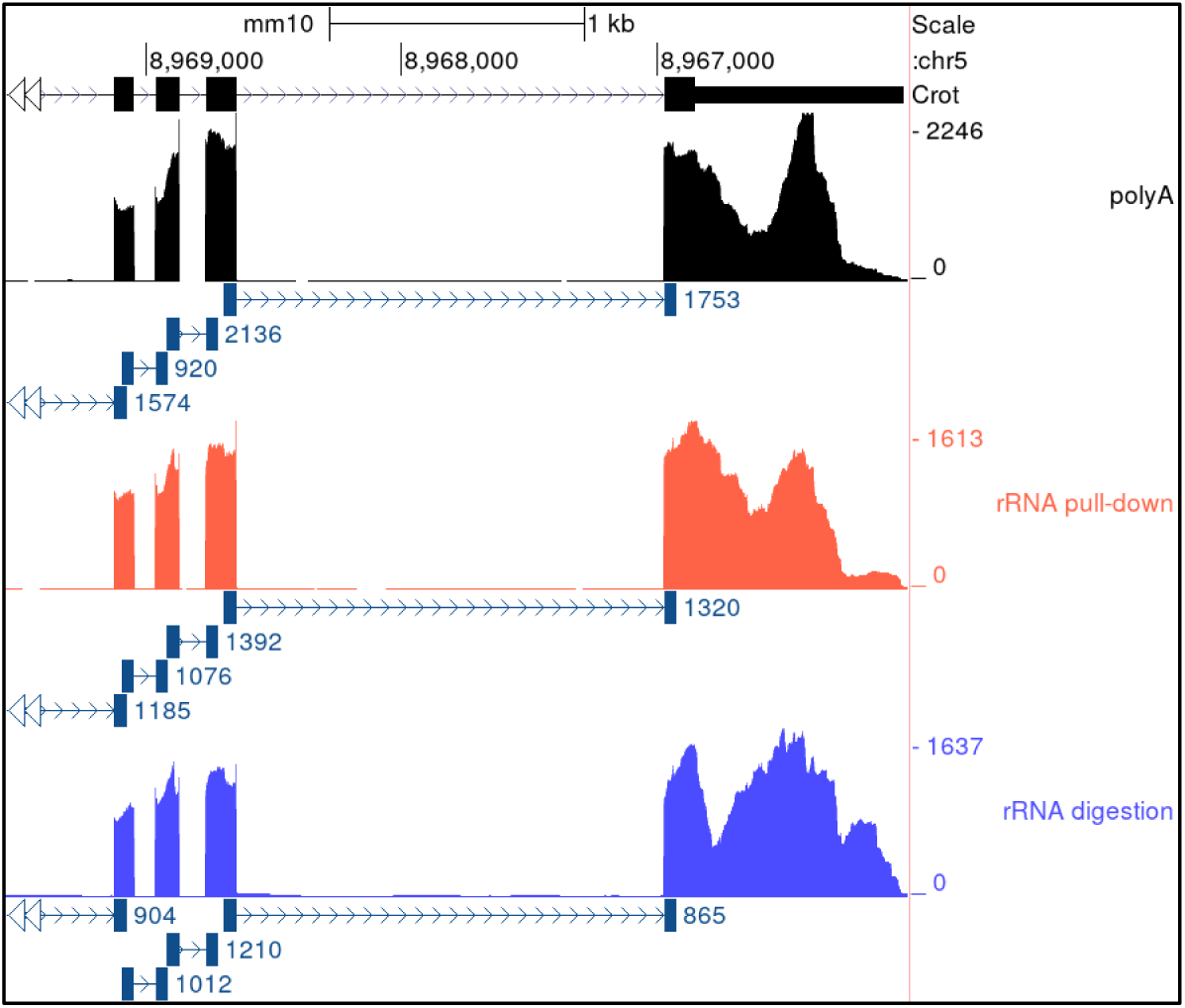
Single-cell and bulk SMART-seq. Coverage of three genes in four individual cells as well as a bulk RNA sample, all prepared by SMART-Seq. SMART-Seq differs significantly from most RNA-Seq protocols considered in this paper, particularly by performing reverse-transcription on full-length transcripts and then performing fragmentation by tagmentation. Data set SMART used.

**Figure 5.**
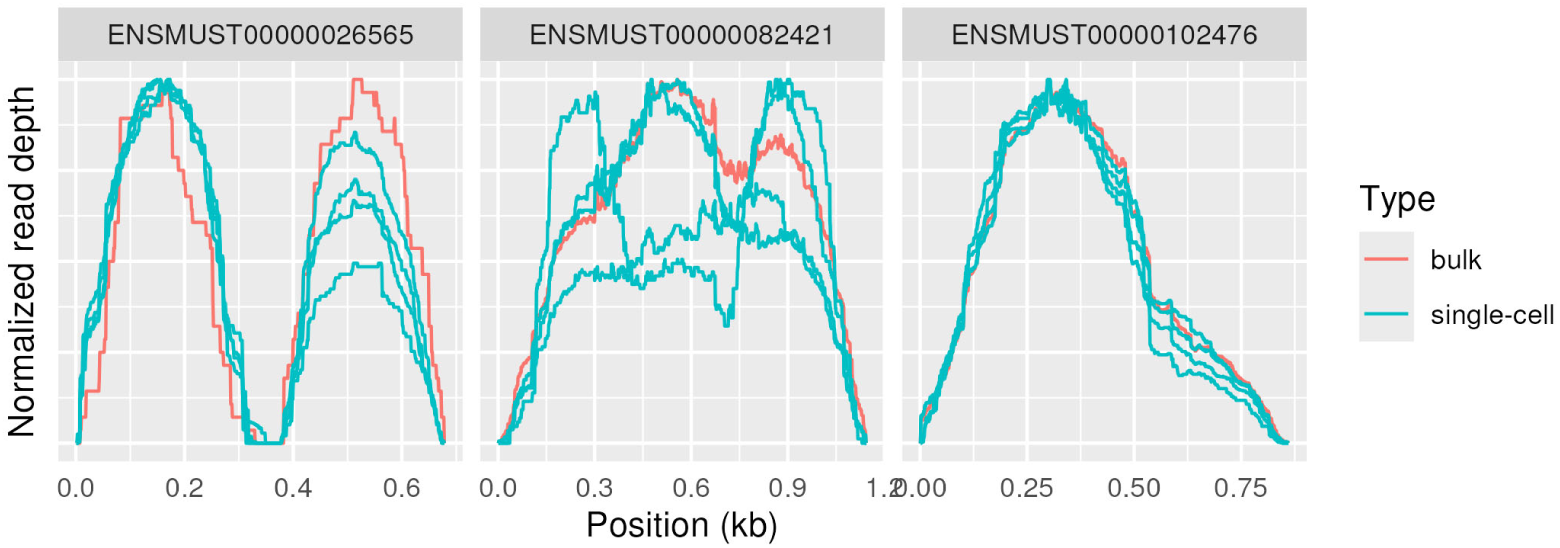
PolyA, rRNA pull-down, and rRNA digestion. Three samples from data set NS1 with varying ribosomal depletion method: polyA selection (top), rRNA pull-down (middle), rRNA digestion (bottom). Coverage is shown of four exons in one gene (Crot). Junction tracks shown as lines with arrows wherever junctions in reads occurred.

While bulk RNA-Seq is our focus, similar non-uniformity is present in single-cell RNA-Seq: samples prepared with a SMART-Seq pipeline from both bulk and individual cells yielded comparable coverage profiles, see Figure 5. Coverage patterns did vary between individual cells in some genes but were mostly consistent.

We will now describe the evidence to address each of the eight putative factors in turn.

### Putative Factor #1: Input RNA sample

One potential explanation for the major hills and valleys is that they could represent accurate abundances of local stretches of RNA in the input sample.

If that were the cause, then polyA selected data should display solely the 3’ bias and none of the other peaks, since the polyA tail is on the 3’ side so a polyA selection assay cannot sequence beyond a gap in the molecule. However, we observe consistent valleys in both polyA and rRNA pull-down libraries, see Figure 4. Valleys must therefore not exist in the input RNA sample. The major hills and valleys therefore do not represent the true nature of the sample.

To investigate sample quality, Figure 6 shows coverage plots for the same sample, which was subjected to greater and greater degradation, as indicated by the RIN scores^1^). As expected, the 3’ bias increases dramatically with degradation. However, the same peaks and valleys are present in all assays, and non-uniformity was not significantly different from constant, see Figure 6 b. This again argues that polyA selection introduces 3’ bias but that the remaining hills and valleys are not due to sample degradation.

**Figure 6.**
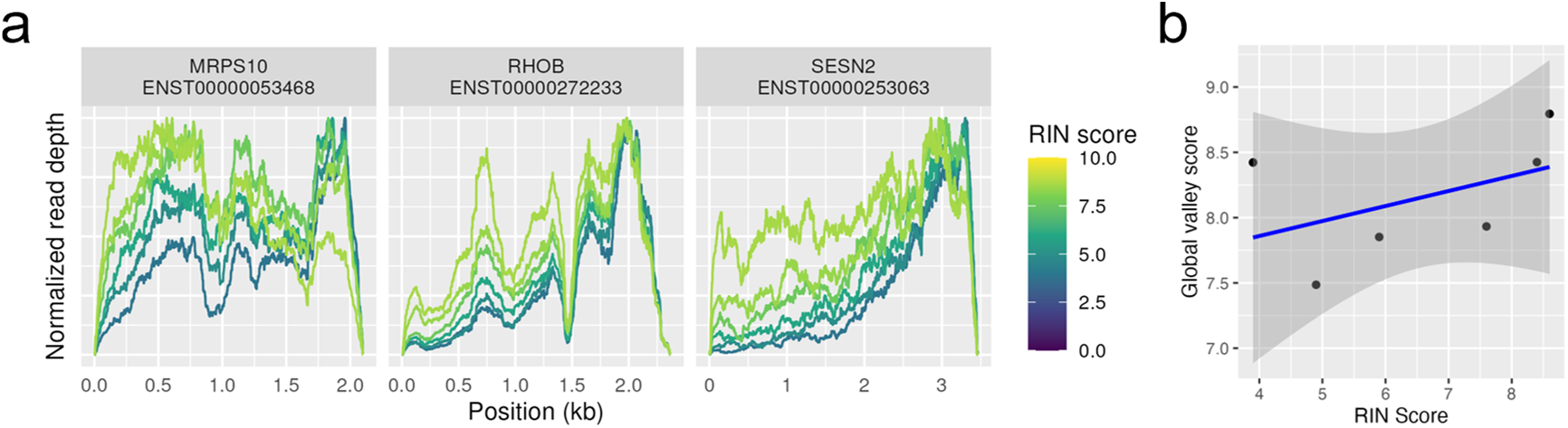
Effect of degradation. Samples from DGRD data set, using SRP073267^1^. These Universal Human Reference samples were subjected to varying amounts of chemical degradation, resulting in a variety of RIN scores. (a) Coverage of three high-expressed single-isoform transcripts are shown. (b) Global valley scores give a measure of non-uniformity, with lower values being more uniform. Coverage was corrected for 3’ bias due to polyA selection before computing the global valley score.

### Putative Factor #2: Fragment sampling variability

We use fragment sampling variability to refer to the variability inherent in the random sampling of which fragments get sequenced. At some level RNA-Seq is modelled as repeatedly selecting colored balls from an urn. In this case, tens of thousands of colors (genes) millions of balls (RNA fragments in the sample) and millions of selections made (sequenced fragments). However, the similarity between technical replicates (Figure 2) demonstrates that the observed non-uniformity is not substantially driven by fragment sampling variability in high-expressed genes. This has previously also been established^17^ by demonstrating considerably lower variability in idealized simulations than is observed in real samples at the same read depth.

### Putative Factor #3: Random hexamer priming

During reverse transcription (RT), the reaction is primed by hybridizing random hexamers to the molecules. This process creates a well-established bias towards specific base pairs at the start and ends of fragments^16^, likely due to a non-uniform distribution of hexamer sequences and variable hybridization affinity. We hypothesize that these biases drive the smallest-scale variations in coverage. To test this, we created position-weight matrices (PWM) of the empirical base composition at the starts and ends of reads for each sample. The PWM-derived score combined from forward and reverse reads predicts the delta from one base to the next in read coverage Figure 7, with the median gene having 0.32-0.51 Pearson correlation with coverage delta across 100 genes in 10 samples, and all genes reached *p* < 0.05 in every sample compared to a randomly permuted PWM. Therefore, we conclude hexamer priming is a primary factor determining small-scale variations in coverage.

**Figure 7.**
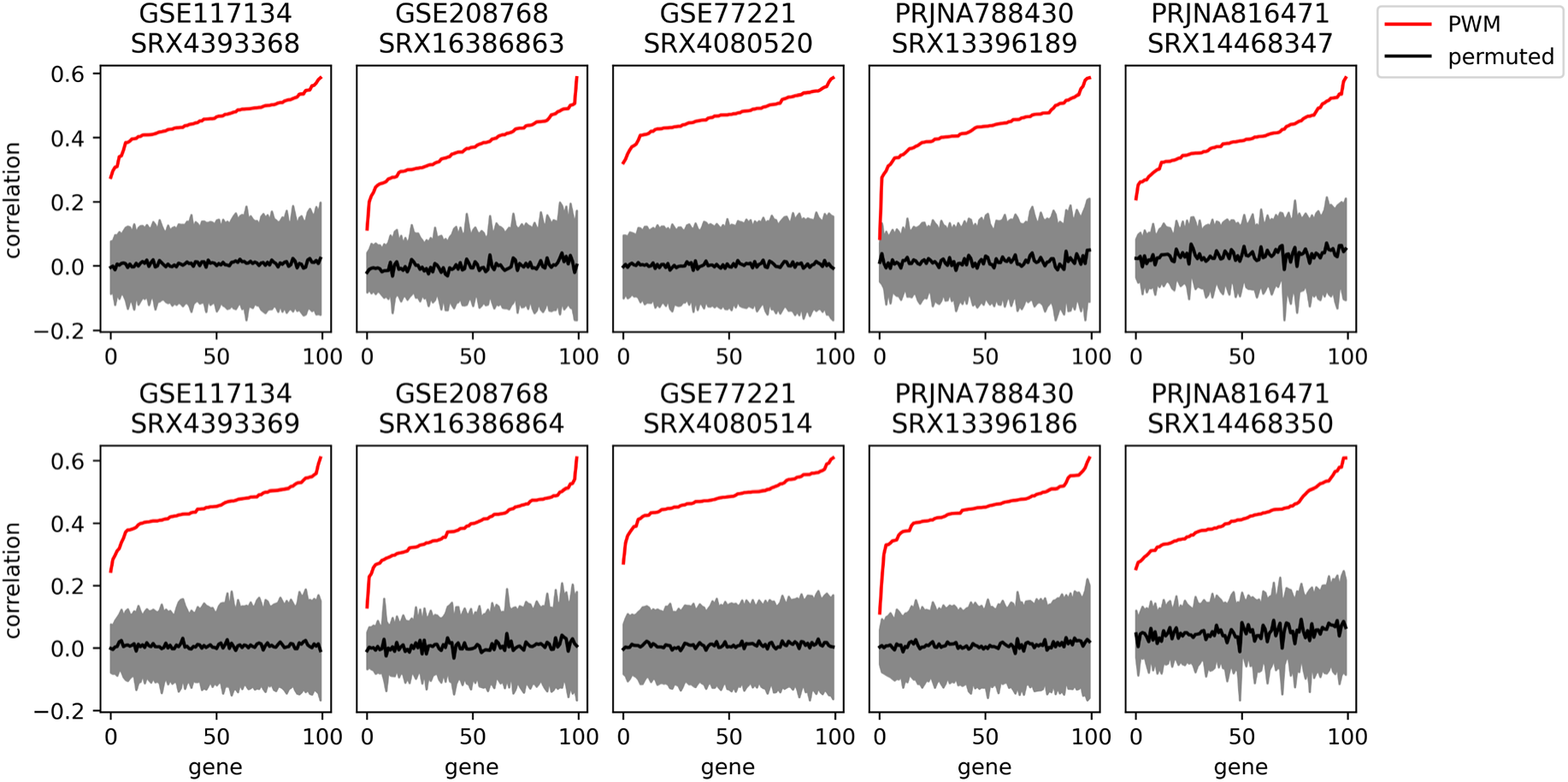
Hexamer binding PWM. We computed an empirical position weight matrix (PWM) for forward and reverse reads of each of ten mouse liver samples from five studies of the GEO data set. PWM scores estimating the propensity for reads to start or end at each base were computed and compared by Pearson correlation to the observed delta in coverage between adjacent positions. Pearson correlations for 100 randomly permuted PWMs were also computed, and all 100 genes were significant at a *p* < 0.05 level in every sample (95% confidence interval shown in gray, mean permuted PWM correlation in black). Data set GEO used.

### Putative Factor #4: Ribosomal depletion

Illumina supports three major strategies of rRNA depletion: polyA selection, rRNA pull-down, and rRNA digestion. Non-specific ribosomal depletion has been proposed as the major driver of non-uniformity^17^, since probes designed to target rRNA might bind non-specifically to other gene sequences. However, non-uniformity is similar across NoSelect, polyA, rRNA pull-down, and rRNA digestion assays, see Figure 4 and Figure 8.

**Figure 8.**
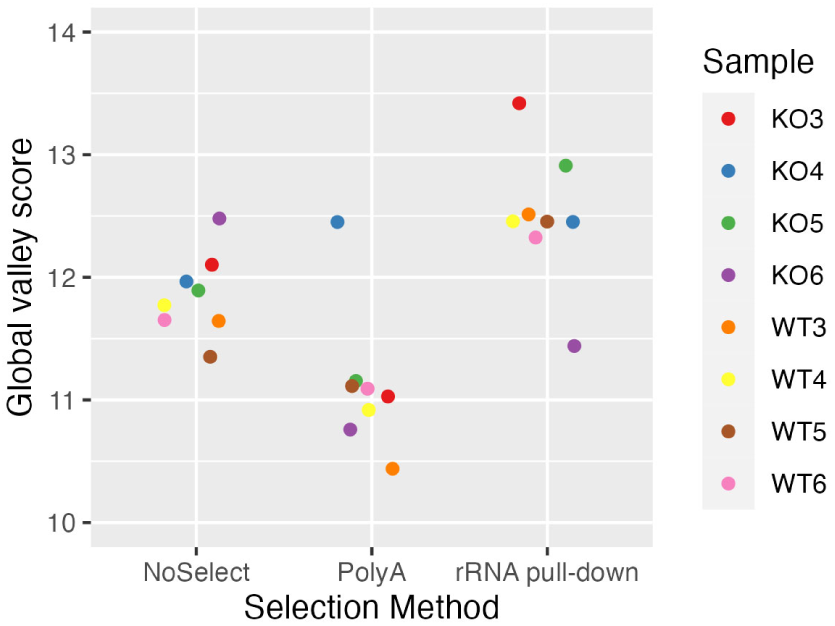
Global valley scores by rRNA depletion. Global valley scores for biological replicates in data set NS2, which were prepared under three different ribosomal depletion protocols (NoSelect, polyA, and rRNA pull-down). Higher values indicate higher non-uniformity.

Figure 4 additionally shows an example of how the three approaches result in somewhat different patterns of coverage, particularly the rRNA digestion.

To assess the true effect of ribosomal depletion, we ran pull-down and digestion protocols both with and without performing the rRNA depletion step to create data set NS1 comparing libraries without any ribosomal depletion (called NoSelect) to traditional rRNA pull-down, rRNA digestion, and polyA kits. NoSelect and rRNA pull-down had comparable coverage profiles, see Figure 9. The rRNA digestion library appeared more different, suggesting that rRNA pull-down has little impact on coverage while rRNA digestion could have an impact. However, the rRNA digestion kit that we used also differed from the polyA, rRNA pull-down, and NoSelect libraries in later steps, so the difference cannot be isolated to just the rRNA depletion step.

**Figure 9.**
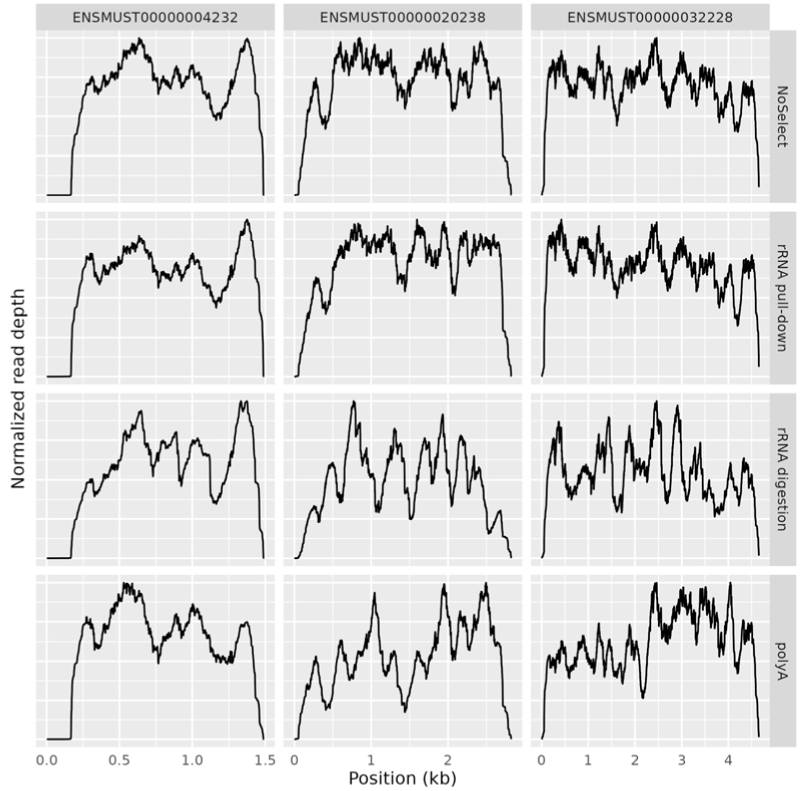
Ribosomal RNA depletion method. Comparison of four samples that varied in rRNA depletion. Data set NS1 used. The rRNA digestion protocol also differs from the others in later steps.

### Putative Factor #5: Fragmentation

Fragmentation generally occurs immediately after ribosomal depletion. If fragmentation were highly enriched in a locus, then it could generate fragments too short to pass the size selection step, resulting in a drop in coverage at that location.

Were this happening, then if we vary the duration of the fragmentation step, we expect to see corresponding variation in the depth of the valleys. We performed this experiment in data set NS1 which was fragmented with divalent cations at elevated temperature. The length of time was varied between 2 minutes, 8 minutes, and 12 minutes, resulting in long, medium, and short fragments, respectively. The initial intention was to test the hypothesis that RNA secondary structure was interfering with the library preparation. The reasoning is that longer fragments would retain more secondary structure and, therefore, should have greater variability of coverage. We, in fact, observed the opposite; see Figure 10. This, instead, is evidence of fragmentation bias causing valleys as hypothesized above.

**Figure 10.**
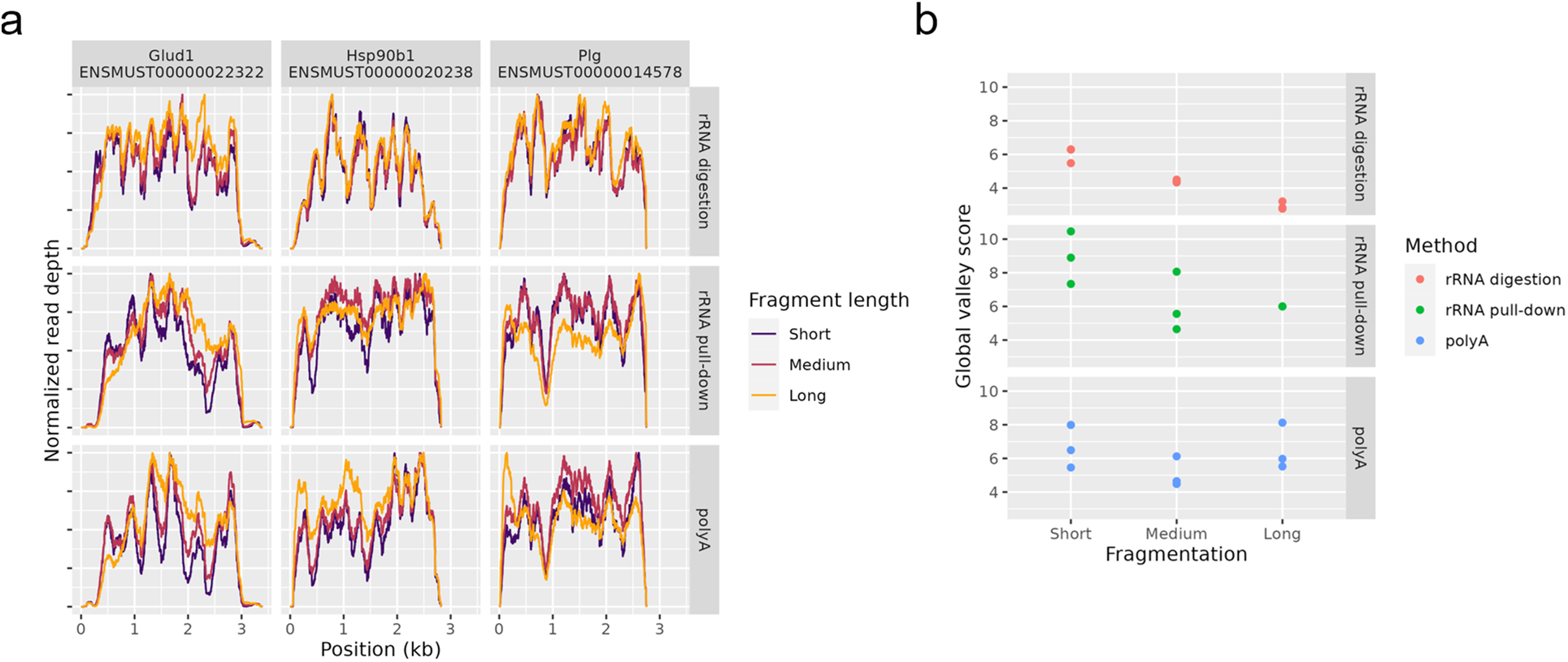
Effect of fragment length. Coverage by different fragment lengths from data set NS1. Fragmentation time was set to either 2 minutes (long fragments), 8 minutes (medium fragments), or 12 minutes (short fragments). (a) Three high-expressed, single

Longer fragments naturally have a smoothing effect on coverage due to how each read pair increases coverage along the entire length of the fragment, which could also explain the observed increase in uniformity at larger fragment lengths.

### Putative Factor #6: Reverse transcription

In our experience, the reverse transcription (RT) step is the most commonly suggested source of artifactual coverage variation, with most believing it is the singular culprit, but it is relatively understudied^18^. Mechanistic explanations typically involve RNA secondary structure interfering with the RT process, since RT is typically performed at lower temperatures (42°C) than PCR (65°C) and other steps. For example, RT can skip over stem-loops, creating chimeric molecules^21^. If this were driving non-uniformity, we should observe reads splicing across gaps where the valleys are located. We do not observe this; splice junctions on reads almost always cross short indels or annotated introns as expected. See Figure 4, which includes a track showing where the spliced reads cross gaps. There are no spliced reads where the valleys are observed. One could argue that these chimeric fragments would be size-selected and that is why we don’t observe the spliced reads. Were that the case, however, we would still expect some chimeric fragments to get through size-selection, so we would still expect to see some splicing at the valleys. But we basically see none.

Additionally, were this the case then we would also expect it to be more severe on longer fragments, since longer fragments can have more secondary structure than short fragments. We observed the opposite: shorter fragments resulted in deeper valleys, see Figure 10.

To investigate the impact of secondary structure further, we attempted to fit coverage by including algorithmically inferred secondary structure as a predictor in the Alpine^5^ model of coverage. We used RNAfold^22^ version 2.4.9 to computationally predict the RNA secondary structure of RNA fragments at 42°C. The amount of secondary structure of each fragment was summarized using the computed mean free energy. Due to computational demands, the starts and end positions of the RNA fragments were rounded to the nearest multiple of 25 bases. Running Alpine with this extra predictor on data set GEO resulted in no distinguishable difference in fit values compared to those of a model without the secondary structure value (data not shown). This suggests that either RNA secondary structure is not a major predictor of non-uniform coverage or that computationally derived estimates of secondary structure are not sufficiently accurate.

We tested this hypothesis in one final way by examining data generated at higher temperatures using thermostable reverse transcriptases, which could alleviate RT-specific non-uniformity. The TGIRT-seq method^23^ incorporates such a thermostable reverse transcriptase. We examined TGIRT-seq data^23^ from the rRNA-depleted universal human reference^24^ samples with reverse transcription performed at 60°C from SRA accession SRP168562. However, these samples still exhibit significant non-uniformity of coverage; see Figure 11, which suggests that much of the non-uniformity comes from steps other than RT. A previous study compared different reverse transcriptases in single-cell RNA-Seq but did not examine coverage non-uniformity and did not make the data publicly available^25^.

**Figure 11.**
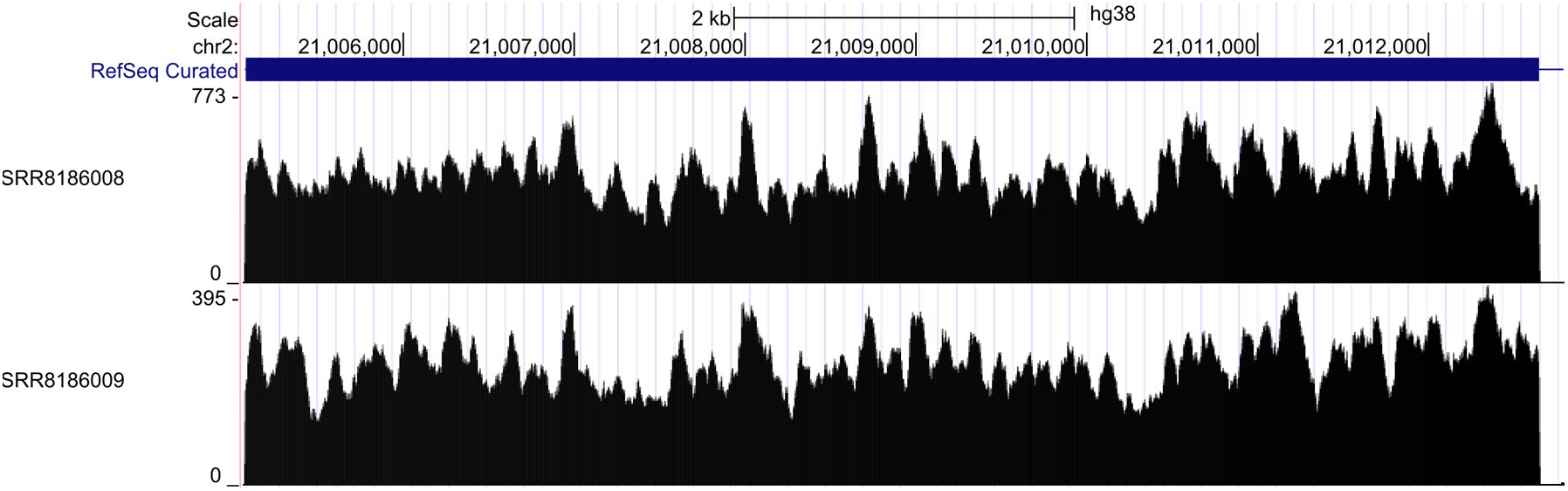
High-temperature reverse transcription. Two technical replicates from the TGIRT data set, which consist of TGIRT-seq data on the universal human reference RNA sample run using high temperature reverse transcription (60°C) from SRA accession SRP168562. Coverage of one exon of the Apob gene is shown

### Putative Factor #7: PCR

After reverse transcription, cDNA fragments are amplified through typically 13 rounds of PCR, which introduces PCR duplicates. These can then be accurately removed by using barcodes, called unique molecular identifiers (UMI)^26^.

It is known^27^ that the molecular makeup of some fragments causes them to amplify with higher efficiency than other fragments. If some fragments fail to amplify well, they will be selected with less frequency to be sequenced which could result in hills and valleys.

We expect that higher PCR efficiency of a particular fragment will be reflected in PCR duplicate rates. This is because higher PCR efficiency of a particular fragment would lead to both a higher chance that at least one PCR copy of that fragment is sequenced and a higher chance that multiple of its PCR copies would be sequenced. Therefore, we can use the duplication rate as a surrogate for amplification efficiency and we can exploit UMIs to determine whether the PCR step is contributing to the non-uniform coverage. If regions of higher PCR duplication are correlated with the peaks, that argues for PCR involvement.

We computed the PCR dupe rate at each position by identifying reads, after deduplication, that overlapped that position, and then computing the fraction of those reads which had at least one PCR duplicate read.

#### Non-uniformity corresponds to PCR efficiency

We observe that in polyA data from data set NS1, the PCR dupe rate does in fact vary substantially from base to base and that this variation does correspond closely to the non-uniformity of (deduplicated) coverage, see Figure 12 a. This relationship was still present, but less clear, in the NoSelect samples that had lower overall PCR dupe rate, see Figure 12 b and Figure S 1. Specifically, we computed the Spearman correlation of smoothed coverage and PCR dupe rate after taking the differences in adjacent bases. This captures whether PCR dupe rate and coverage both tend to move in the same direction. Across the chosen 100 genes, both samples had non-zero correlation (1-sample Wilcoxon test, *p* < 10^-9^). The overall higher PCR dupe rates observed in polyA samples compared to NoSelect samples could be driven by the lower diversity of molecules after polyA selection.

**Figure 12.**
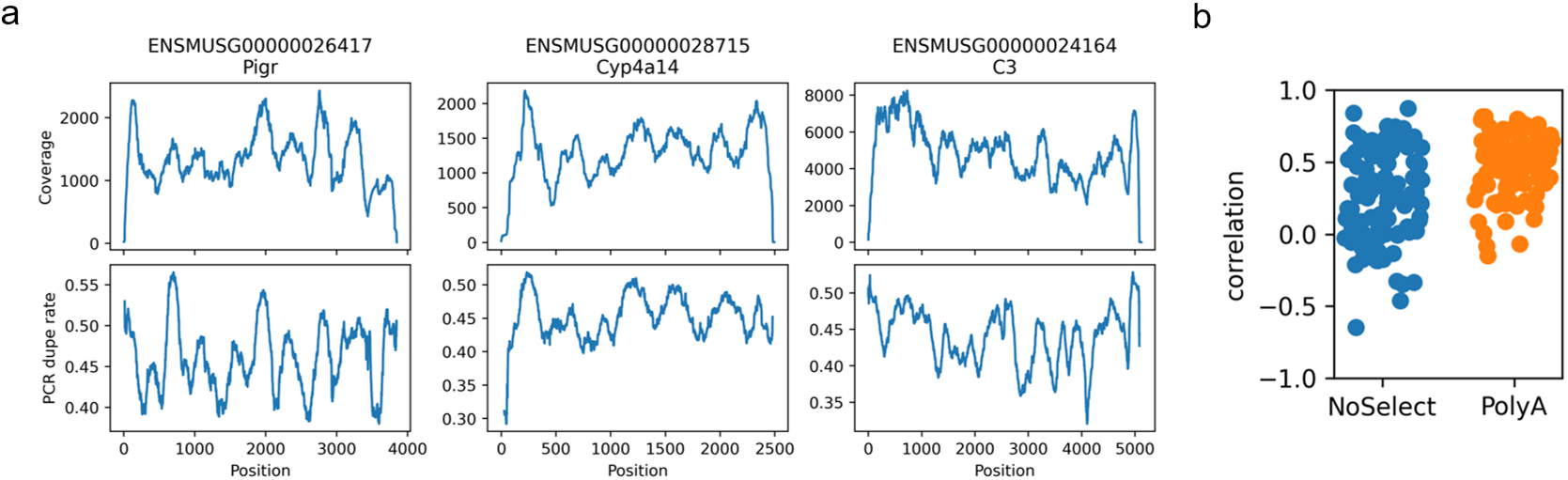
PCR dupe rate by position. (a) In a polyA-selected mouse liver sample from data set NS1, coverage non-uniformity (top row) and PCR dupe rate (bottom row) in three select genes. We use PCR dupe rate as a proxy for PCR efficiency, since fragments that are more efficiently copied are more likely to result in multiple sequenced copies. Plotted coverage is after removal of PCR duplicates. Ribosomal reads were computationally removed prior to computing duplicates. (b) Spearman correlation between PCR dupe rate and deduped coverage, after smoothing and taking the difference of adjacent bases in each. Correlations in 100 single-isoform genes are shown in two samples with high PCR dupe rate (PolyA) and low PCR dupe rate (NoSelect). Both are statistically non-zero (1-sample Wilcoxon test *p* < 10^-9^).

#### GC content and PCR amplification

It is known that GC content affects PCR amplification^28^, and using UMIs to compute PCR dupe rate allows us to separate out GC effects in PCR amplification versus GC effects in other aspects of the library preparation and sequencing. We find that PCR dupe rates depend upon GC content in a library-specific manner, see Figure 13.

**Figure 13.**
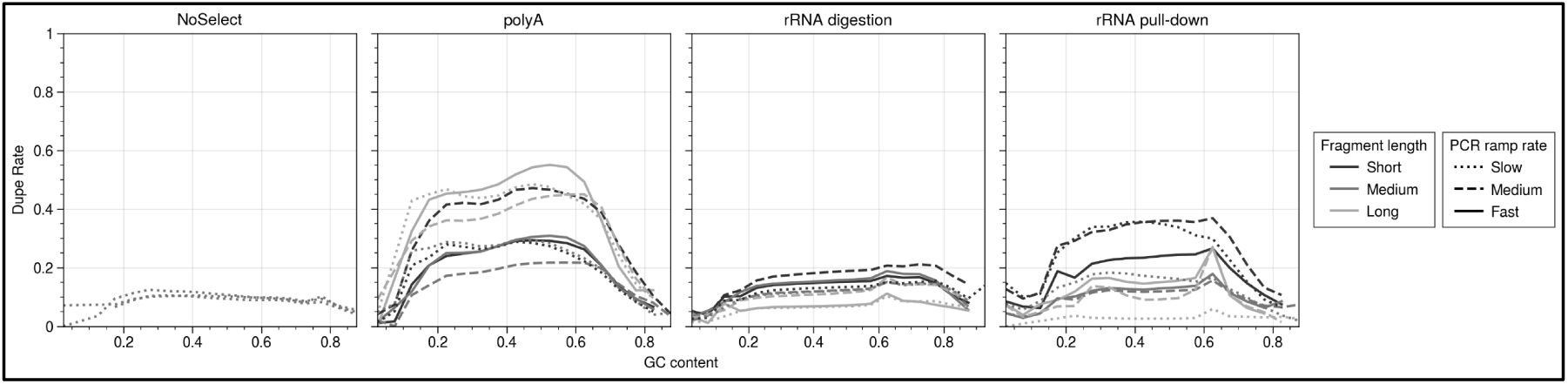
PCR duplicate rate by GC content. PCR dupe rate is determined by use of unique molecular identifiers. Dupe rate in each sample is shown versus read GC content. Data from data set NS1 was used. Overall dupe rate varies substantially between rRNA depletion methods, likely due to changes in library complexity after selection.

#### Effect of PCR cycle number

We hypothesized that if PCR was driving non-uniformity of coverage, then using more PCR cycles would lead to larger non-uniformity. To test this, we used mouse testis data from a study^29^ that varied PCR cycle numbers from 8 to 13 and used UMI tags. Looking in highly expressed genes, we observed no substantial trends in either non-uniformity of coverage itself or in the non-uniformity of the PCR dupe rate as the PCR cycle number increased, see Figure 14. Overall PCR dupe rates do increase as PCR cycle numbers increase, though this is also a function of the corresponding decrease in input RNA that were used for the higher PCR cycle numbers, see Figure S 2.

**Figure 14.**
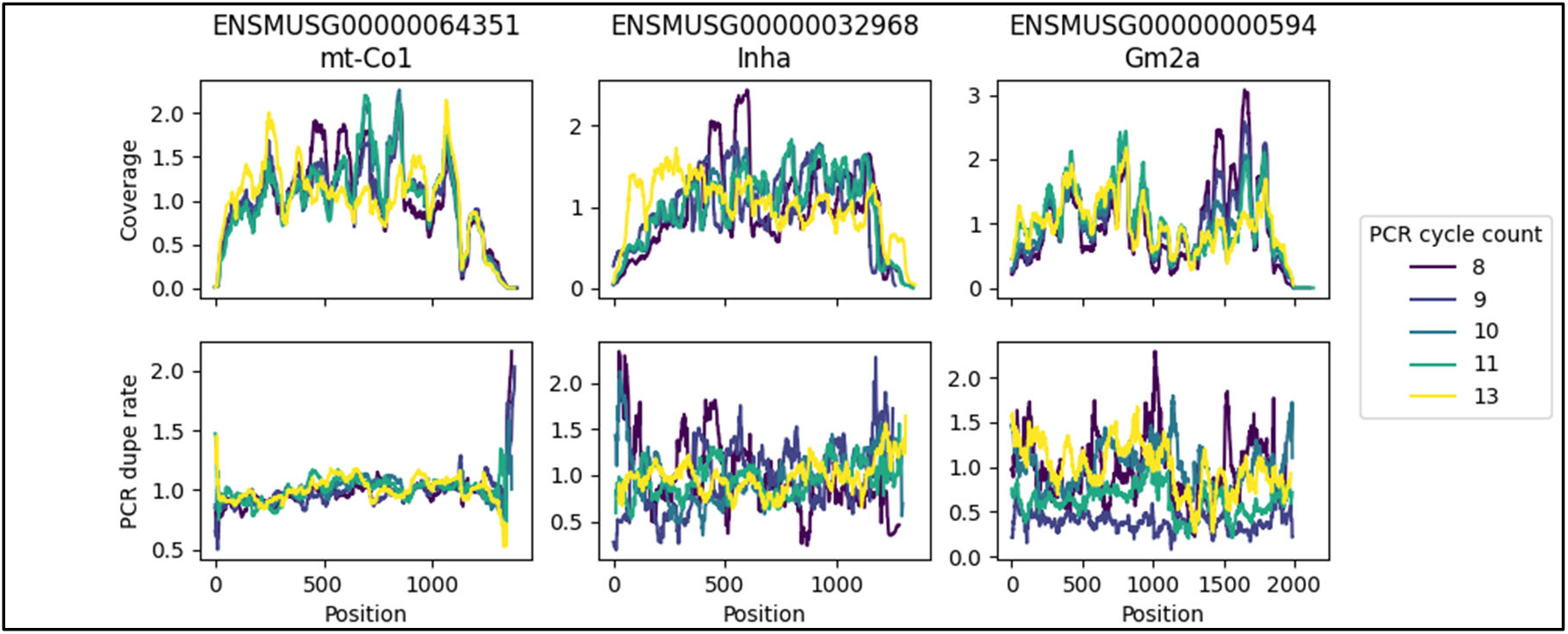
PCR cycle counts - relative values. Mouse testis data from SRA accession PRJNA416930 with varying PCR cycle numbers. Normalized coverage and normalized PCR dupe rate are plotted, due to overall PCR dupe rate depending highly on PCR cycle numbers. PCR dupe rate is determined by UMI tag and is computed as the fraction of deduplicated reads that originally had at least one duplicate. Samples with higher PCR cycle numbers also had lower input material. Coverage is computed after removal of PCR duplicates.

#### PCR ramp rate

The ramp rate used by the thermocycler during PCR has been suggested to control non-uniformity^19^. Data set NS1 has varying PCR ramp rate, but we observed no effect on non-uniformity, see Figure 15.

### Putative Factor #8: Alignment

Non-uniformity could be due to regions that are difficult to align due to either multi-mapping or failing alignment entirely. We extracted 100bp transcriptomic sequences as synthetic ‘reads’ around each identified peak and valley in one sample from data set NS1 and used STAR to align them to the genome.

**Figure 15.**
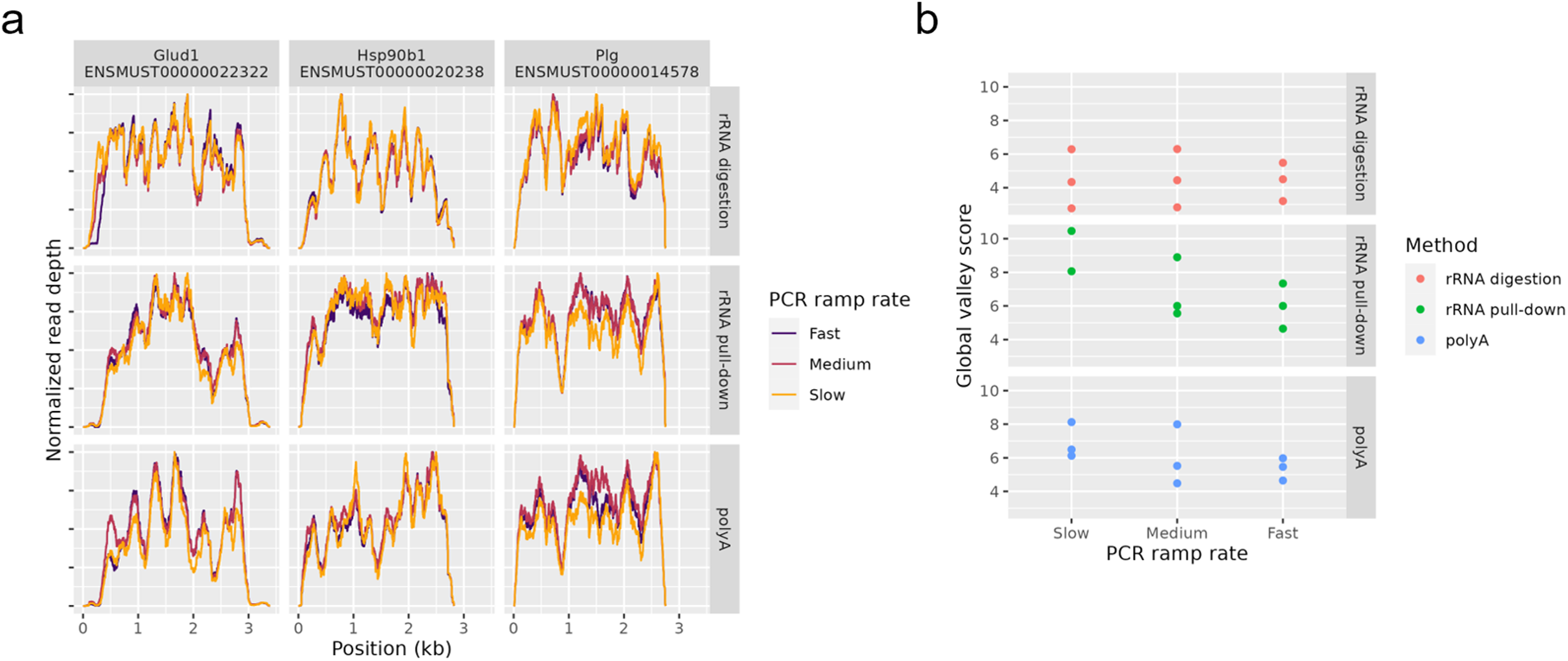
PCR ramp rate. Coverage by different PCR ramp rates from data in data set NS1. PCR ramp rate is the rate at which the thermocycler changes temperature. (a) Coverage of three top expressed genes. (b) Global valley scores for each sample. Higher values indicate more non-uniformity.

All reads aligned successfully, but alignments were unique for 96.5% (805/834) of reads from peaks and for 93.9% (493/525) of reads from valleys, for a *p*-value of 0.017 (Fisher’s exact test), indicating a small increase in multi-mapping in valleys.

This implies that alignment issues play some role, but it is not clear from this analysis how much of a role. To assess that, coverage plots were computed using only uniquely mapping reads or including multi-mapping reads. If multi-mapping regions were driving valleys, we would expect this comparison to show differences in valleys. Indeed, some genes show clear patterns where valleys in unique mapping coverage are filled in when including multi-mappers, see Figure 16 left column. However, switching to always including multimapping reads would also affect other genes, see Figure 16 rightmost column, where multimapping reads appear more likely to originate from other loci. However, most genes show relatively few multi-mappers; 72 of the 100 genes have under 10% multi-mapping reads at every base of coverage in all samples of the GEO data set. We conclude that alignment issues play some role but only in specific regions.

**Figure 16.**
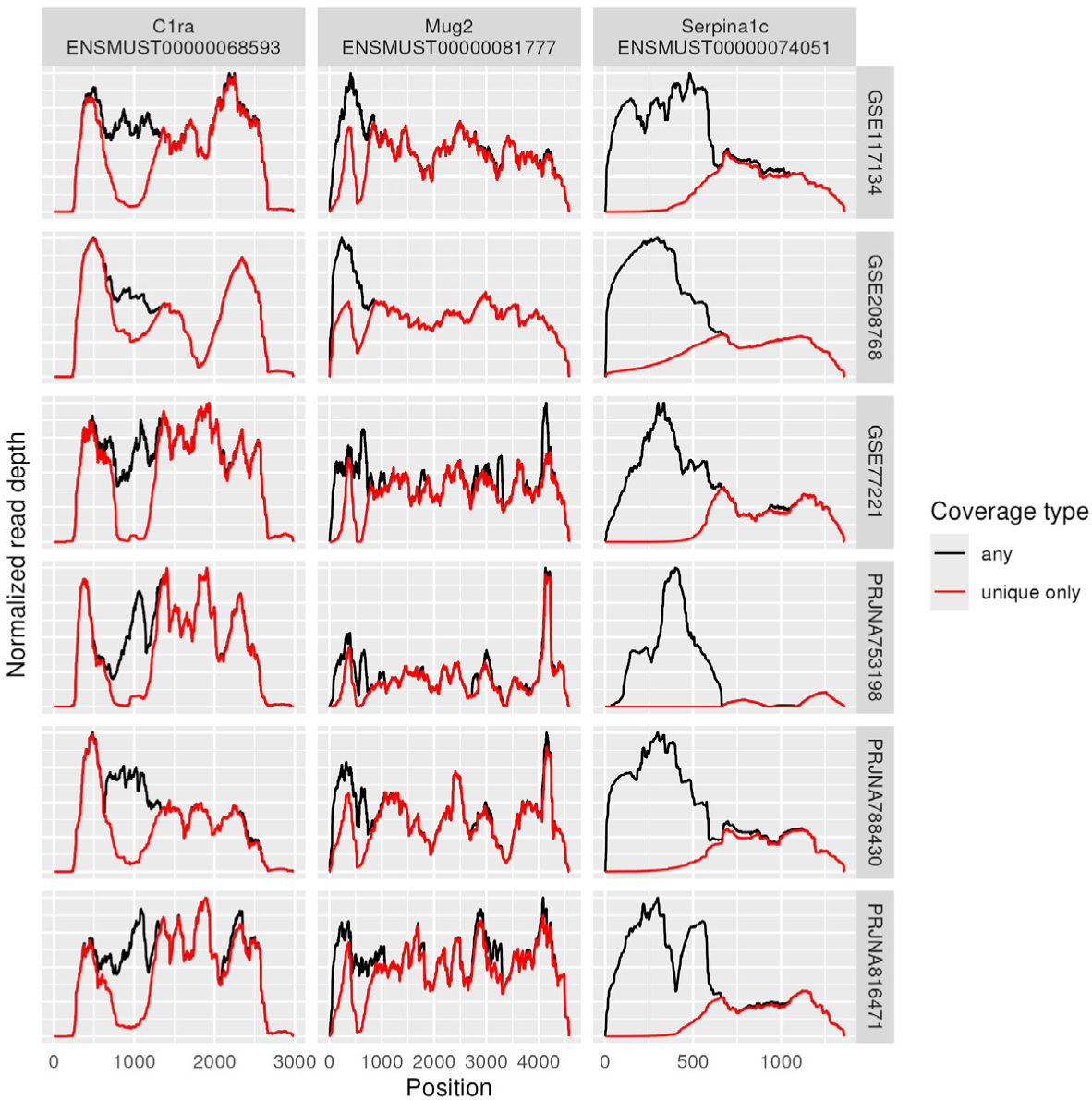
Multi-mappers. Coverage plots computed for just unique mappers as well as for all mappers (including multi-mappers). Three genes chosen for having the largest divergences in coverage between including multi-mappers or not. While some genes (leftmost column) show clear valleys that are filled in by inclusion of multi-mappers, others (rightmost column) seem more likely to have multi-mappers that truly originated from a different locus. Data set GEO used.

### Computational models of coverage non-uniformity

Lastly, we consider whether current computational methods can model coverage as a function of genomic sequence. Alpine^5^ models the rate of sequencing each potential fragment in a gene as a function of GC content, position, and sequence near the start and end locations. Non-uniformity of coverage was incompletely captured by the Alpine models, with varying results across different genes and different studies, see Figure S 3.

## Discussion

Understanding coverage bias in RNA-Seq turns out to be a difficult problem. Our goal here was to clarify the landscape of current progress.

Sources of non-uniformity in RNA-Seq remain incompletely characterized. We dispensed with several commonly proposed mechanisms by demonstrating they do not explain much of the variability, if at all. First, shorter fragments did not improve uniformity despite the expected lower propensity for secondary structure in shorter fragments. Indeed, longer fragments gave somewhat more uniform coverage.

Second, lowering PCR temperature ramp rate did not substantially alter coverage. Third, omitting ribosomal depletion yielded remarkably similar coverage to the original rRNA pull-down library preparation protocol. Fourth, decreasing the number of PCR cycles used did not improve uniformity. Lastly, while we did not systematically test the reverse transcription step, public data using a high-temperature reverse transcriptase does not appear to have substantially improved uniformity.

We observed that Alpine, a model of RNA-seq coverage, leaves a sizeable portion of the non-uniformity unexplained. Its performance varied from sample to sample and gene to gene, indicating that factors outside of GC content, start and end sequence, and position within the transcript contribute to the non-uniformity. Including a measure of inferred secondary structure content did not improve the Alpine performance, suggesting that either secondary structure could be less important than expected or that computational methods provide an inadequate estimate of it. This was surprising as secondary structure interference in reverse transcription was often suggested as the primary source of non-uniformity.

It remains challenging to assess the impact of any specific part of the library preparation pipeline due to the combination of all parts. For example, the difficulty in assessing fragmentation bias in RNA-Seq is that, because of the random hexamer step, the reads produced rarely represent the true ends of the fragments. Short read RNA-Seq protocols have been designed to sequence to the very end of fragments, so conceivably this approach can be used in typical RNA-Seq to be able to assess fragment ends. This experiment will be the focus of a future publication. Moreover, bulk RNA-Seq is an average signal over many cells, each of which is a snapshot in time when RNA is actively being both transcribed and processed and degraded. We observe only a combined effect of every cell being somewhat different, and post-translational modifications to RNA such as methylation could impact sequencing without being readily assessable. However, the non-uniformity issue is basic, and a resolution will be impactful.

Another difficulty is the observed variability between studies, which may explain why this topic has proven difficult to reproduce. Specifically, this could be why we did not replicate results about the rRNA depletion step, PCR ramp rate, or fragment length.

Despite these difficulties, several things are clear. First, polyA-selection induces the expected 3’ bias in longer transcripts. Second, efficiency of PCR correlates well with non-uniformity at least in samples with higher PCR duplicate rate. This makes it plausible, if not likely, that PCR drives part of the non-uniformity. PCR efficiency also had a GC content-dependent effect. However, since reducing PCR cycle numbers did not improve non-uniformity, PCR efficiency may be merely acting as a proxy for efficiency at some other step, particularly reverse transcription. Third, non-uniformity appears to have multiple sources, and addressing any one or two of these is not sufficient to remove all non-uniformity. Fourth, non-uniformity is highly consistent across technical replicates, reasonably consistent across biological replicates, and only modestly consistent across studies. Despite that, some peaks and valleys appear to be consistent across most studies, suggesting that it is a feature of the sequence itself that causes those regions of non-uniformity.

We demonstrated that the NoSelect protocols, which omit rRNA depletion, are viable choices given rapidly decreasing sequencing costs. However, NoSelect produced very similar coverage to the rRNA pull-down kit it was based on and had similar genomic content (e.g., exonic, intronic, intergenic) excluding rRNA, so for most purposes, the rRNA pull-down kit (Ribo-Zero) would be equivalent.

We observed no effect of PCR ramp rate on coverage despite a previous report of its impact^19^. This may be explained by the fact that all three ramp rates were performed by adjusting the ramp rate of the same PCR machine, while the prior study used different machines to achieve different ramp rates.

In conclusion, the non-uniformity of coverage in RNA-Seq continues to defy simple explanations. Further work is needed on multiple fronts: computationally accounting for or correcting non-uniformity, identifying the largest contributors to non-uniformity, and improving experimental protocols to avoid non-uniformity. Improvements to any of these areas could substantially affect RNA-Seq downstream analysis and its promise of accurate isoform level analyses.

## Methods

### Animal care, tissue collection, and RNA extraction

Wild-type, twelve-week-old male C57BL/6J mice were bred in-house at the animal facility of the University of Pennsylvania. The mice were euthanized through carbon dioxide induced asphyxiation and liver, heart or lung samples were dissected and snap-frozen in liquid nitrogen. RNA was extracted from the tissue using TRIzol (ThermoFisher Scientific, catalog no. 15596018) and RNeasy Mini Kit (Qiagen, catalog no. 74104), according to manufacturers’ protocols. After extraction, total RNA was analyzed on a BioAnalyzer 2100 (Agilent) to check for integrity. All procedures were approved and carried out in accordance with the Institutional Animal Care and Use Committee of the University of Pennsylvania.

### Library preparation and sequencing of data set NS1

Except where noted, we constructed all libraries according to the manufacturer’s protocols and used 13 cycles of PCR during the final library amplification step. To allow us to assess duplication rates with UMI sequences, we used the xGen UDI-UMI adapters (IDT, catalog no. 10005903) for all libraries, diluting the initial 15 μM stock of adapters to 3 μM in NGS adapter buffer (10 mM Tris pH 8.0, 0.1 mM EDTA, 100 mM NaCl).

We used the TruSeq Stranded mRNA library preparation kit (Illumina, catalog no. 20020594) and the TruSeq Stranded Total RNA Human/Mouse/Rat library preparation kit (Illumina, catalog no. 20020596) to prepare the PolyA and rRNA pull-down libraries, respectively. For the rRNA digestion libraries, we used the Illumina Stranded Total RNA Prep Ligation with Ribo-Zero Plus kit (Illumina, catalog no. 20040525). This kit uses a different strategy than the TruSeq kits for ligating indexed adapters to cDNA fragments that is not compatible with the xGen adapters. In order to use the same adapters, and their included UMIs, across all libraries, we modified the protocol for the Illumina Stranded Total RNA Prep Ligation as follows: 1) During the ‘Ligate Adapters’ step, replace the RNA Index Anchors with the diluted xGen adapters, and 2) during the ‘Amplify Library’ step, replace the Index Adapters with the PCR Primer Cocktail (PPC) from the TruSeq Stranded Total RNA kit. We performed all other steps according to the manufacturer’s protocol. All PolyA and ribo-depleted libraries used 200 ng of input RNA.

For the NoSelect libraries, we used 20 ng of total RNA as input into the TruSeq Stranded Total RNA library preparation protocol, skipping the ‘Deplete rRNA’ step and starting with the ‘Fragment and Denature RNA’ step (see supplementary methods for detailed protocol). All remaining steps in the library preparation used reagents from the TruSeq Stranded Total RNA library preparation kit and followed that protocol. Since the majority of reads from the No Select library come from rRNA reads, we prepared and sequenced three No Select libraries from each sample/condition in order to get a sufficiently diverse sampling of non rRNA molecules.

For the experiments where we varied RNA fragmentation, we used the following times for the 94°C incubation during the fragment RNA step: 2 minutes (short), 8 minutes (medium), or 12 minutes (long). Similarly, for experiments where we varied PCR ramp rates, we used the following Thermocycler ramp rate settings during the library amplification step: 3°C/s (fast), 1.5°C/s (medium), or 0.5°C/s (slow).

Libraries were sequenced on an Illumina NovaSeq 6000.

### Library preparation and sequencing of data set NS2

Libraries were prepared and sequenced as described for data set NS1 except that 11 PCR cycles were used, adapters were diluted to 1.5 μM in the NGS adapter buffer, and 15 ng of total RNA was used as input the library preparation.

### NoSelect, polyA, and rRNA pull-down and digestion kits contents

Data set NS1 was performed with varying rRNA depletion techniques, including without any depletion, see Figure S 4. The NoSelect libraries proved successful, with 89-90% of reads being ribosomal, 93% reads aligning uniquely after deduping by UMI tags, see Table 2. With 6 pooled libraries, this gave a total of 781 million reads yielding 30.5 million deduped, non-ribosomal reads in the NoSelect liver sample.

**Table 2.**
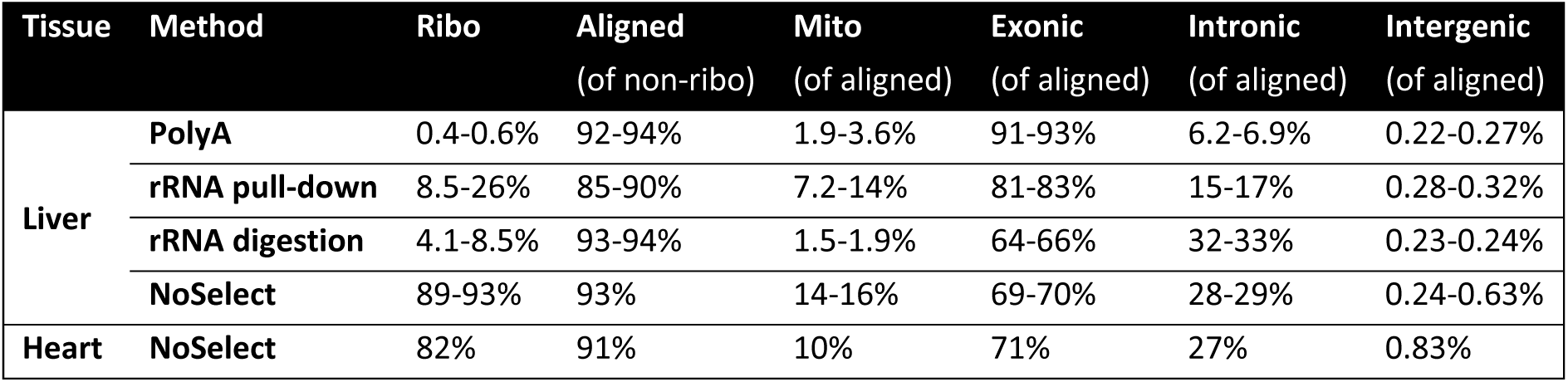
Sequencing content Fraction of reads after deduplication falling into each category for libraries from each mRNA enrichment method. Ranges denote the minimum and maximum values of all the libraries. Ribosomal reads were removed prior to deduplication and alignment via mapping to a ribosomal RNA database using BLAST. Only uniquely aligned reads were counted as aligned and all further columns refer to percentages of uniquely aligned reads.

| Tissue | Method | Ribo | Aligned<br>(of non-ribo) | Mito<br>(of aligned) | Exonic<br>(of aligned) | Intronic<br>(of aligned) | Intergenic<br>(of aligned) |
| --- | --- | --- | --- | --- | --- | --- | --- |
| Liver | PolyA | 0.4-0.6% | 92-94% | 1.9-3.6% | 91-93% | 6.2-6.9% | 0.22-0.27% |
|  | rRNA pull-down | 8.5-26% | 85-90% | 7.2-14% | 81-83% | 15-17% | 0.28-0.32% |
|  | rRNA digestion | 4.1-8.5% | 93-94% | 1.5-1.9% | 64-66% | 32-33% | 0.23-0.24% |
|  | NoSelect | 89-93% | 93% | 14-16% | 69-70% | 28-29% | 0.24-0.63% |
| Heart | NoSelect | 82% | 91% | 10% | 71% | 27% | 0.83% |

PolyA libraries contained the highest fraction of aligned reads and the highest fraction of exonic reads. NoSelect, rRNA pull-down and rRNA digestion libraries showed both much higher intronic content and slightly higher intergenic content than did polyA libraries. Overall content distribution was similar between rRNA pull-down, rRNA digestion and NoSelect, after removing ribosomal reads.

### Bioinformatics analysis

Sequenced libraries were aligned with STAR v2.7.6a^30^ and duplicate reads (identified by UMI tags) were dropped with umi_tools dedup v1.1.1^31^. All NoSelect libraries of the same sample were merged for downstream analysis. Results were quantified and normalized within each selection method by a down-sampling method, PORT v0.8.5e (github.com/itmat/Normalization/). PORT also identified and removed rRNA from the samples prior to normalization and quantification; this is done by BLAST^32^ search of rRNA to a prepared sample of ribosomal DNA due to the lack of high-quality annotation of rDNA in reference genomes. A random selection of 500 reads from one sample were processed by hand through the NCBI’s BLASTN web-interface to classify sample origin and read type and cross-reference with PORT-inferred rRNA classification and identified high congruence between the automatic and manual classification.

Alpine^5^ models the rate of sequencing each potential fragment of single-isoform genes with a Poisson GLM. Alpine was applied to a set of 100 single-isoform genes chosen to be high expressed in all liver samples. Alpine was modified to allow for the use of gene length in the model since positional bias in PolyA selected data was observed to be highly length-dependent (long genes exhibit significant 3’ bias but short genes do not). The model included the following factors: fragment GC content; relative position in fragment; relative position interaction with gene length; fragment GC stretches (40bp at 80% GC and 20bp at 80% GC); fragment length; gene; and 21bp-wide variable length Markov model on the sequence on fragment start and end locations. Alpine defaults were used for GC and relative position knots. Fit values were used to estimate the effects of GC content and position on read coverage.

PCR copies were counted by binning reads based on the GC content of their first read (in 5%-wide GC bins) and using the umi tools group command on the reads aligned to chromosome 19.

### Selected transcripts

To avoid the difficulty of distinguishing alternative splice forms of the same gene, we selected for each data set a collection of 100 single-isoform genes. These were chosen from the Ensembl v109 annotation of the human genome and the Ensembl v102 annotation of the mouse genome. Only genes of protein coding or lncRNA biotypes are included and genes were selected that had lengths between 600 and 7000 base pairs to represent typical lengths of protein coding genes. Among these genes, the 100 highest expressed in the data set were selected.

### Coverage plots

Coverage plots can be computed in several ways. All data was aligned to the genome and so includes introns. When possible, we present data from single-isoform genes and so compute coverage relative to position within the transcript, which effectively removes introns. Moreover, in this case, we also know both start and end location of the read and can assume that the entire region between the start and end is covered by the read (i.e., there are no junctions in the transcriptome). So coverage plots are computed by incrementing all positions between start and ends of the paired read. All data sets used paired reads. In data sets with unique molecular identifiers (UMIs), coverage is computed after performing deduplication of PCR duplicates. Normalized coverage is computed by scaling each sample and each gene individually to the range of 0 to 1, i.e., by dividing by the maximum coverage of that sample within that gene. Except where noted otherwise, coverage was computed using only uniquely aligned reads. Data is shown with the 5’ to 3’ ends going from left to right.

### Peak and valley finding

Fragment coverage tracks were first smoothed by a Gaussian kernel of standard deviation 30 bases. Initial valleys were identified by local minima between two local maxima such that the difference in coverage from minima to both maxima was at least 10% of the average coverage of the gene. A “global valley score” was computed as the total amount of water that every valley could contain, normalized by average coverage and transcript length. Larger global valley scores correspond to coverage with larger valleys.

Low expression regions were determined as the portion of valleys that were in the bottom third of coverage. High expression regions were neighborhoods of peaks in the top third of coverage. Random 100 base pair sections of the transcriptome sequence were selected from both low and high expression regions determined from one sample (liver rRNA pull-down, slow PCR ramp rate, medium fragment size) and were aligned with STAR to the genome to assess difficulty of alignment.

### Global valley score

To measure the amount of non-uniformity in a sample, we define the “global valley score”. First, for each identified valley, we compute the amount of “water” that can be contained in that valley: the sum of min(*left*, *right*) – *cov_i_* where *left* and *right* are the coverage at the left and right peaks and *cov_i_* is the coverage at the *i*th base between the peaks. This is the local valley score. The global valley score is the sum over all valleys of the local valley score, normalized by average coverage and transcript length. Larger values indicate more non-uniform coverage.

In order to account for 3’ bias in polyA selected samples, a reference sample was chosen where we expected limited 3’ bisa. For data set NS1, this was the NoSelect sample. For data set DGRD, this was the highest RIN score sample. For each sample, the log ratio of that sample’s coverage and the reference’s coverage was regressed against distance from 3’ end, relative position (i.e. position on the transcript from the 5’ end divided by the length of the transcript), and the interaction term. The fitted line gives an exponential fit of coverage in a sample as a function of position, with the same fit parameters being used for every gene. A corrected coverage was computed by dividing the actual coverage by this fit value, which removes the 3’ bias. Global valley scores were computed using this corrected coverage.

### Hexamer binding and PWM analysis

We computed an empirical position weight matrix (PWM) for forward and reverse reads of each of ten mouse liver samples from five studies. The PWMs used the first and last 12 bases of each fragment, excluding soft-clipped bases. The PWMs were compared to transcript sequences for each of 100 high-expressed single-isoform genes. All ten samples were sequenced such that the reverse read aligned to the transcript and therefore indicated the 5’ end of the fragment. A composite PWM score was computed by subtracting the forward read PWM ending at position k from the reverse read PWM score starting at position k+1. This estimates the difference in the propensity for reads to start at k+1 minus the propensity to end at k and therefore is hypothesized to approximate the change of coverage.

Pearson correlations were then computed with the difference of expression at position k+1 and position k, after removing a smoothed fit of coverage to account for large-scale variations in coverage.

One correlation was computed for each of the 100 genes in each of the 10 samples. To compute a p-value, we computed 100 permutations of each PWM by scrambling the positions within the forward and reverse PWMs, and correlations with the permuted PWMs were computed. This test, therefore, accounts for the dependence of nearby bases in coverage. A one-sided p-value test was used as the correlation with the observed PWM was expected to be positive.

The authors declare no competing interests.

## Data availability

Sequencing data for data sets NS1 and NS2 have been made available from the Gene Expression Omnibus (GEO) at accessions GSE288460 and GSE288461, respectively. All other data were from publicly available data with accessions specified in Table 1.

