## Supplemental Methods and Figures for "Sources of non-uniform coverage in short-read RNA-Seq data"

### Supplemental Methods

#### Modification to TruSeq Library Preparation Protocol for No Select Libraries

This portion of the protocol is adapted from the 'Deplete rRNA' and the 'Fragment and Denature RNA' steps in the August 2019 revision of the "TruSeq Stranded Total RNA With Illumina Ribo-Zero Plus rRNA Depletion" (Document # 1000000092426v01).

<u>Consumables</u>

Nuclease-free water

RNAClean XP beads

80% ethanol (freshly-prepared)

ELB – Elution buffer (from TruSeq Stranded Total RNA library prep kit)

EPH – Elute, Prime, Fragment High Mix (from TruSeq Stranded Total RNA library prep kit)

<u>Preparation</u>

Save the following program on the thermal cycler:

- Choose preheat lid option and set to 105°C
- 94°C for 8 minutes
- Hold at 4°C

<u>Procedure</u>

1. Dilute 20 ng of total RNA in 30 µL of nuclease-free water.
2. Add 60 µL RNAClean XP beads.

3. Pipette up and down until beads are fully resuspended.
4. Incubate at room temperature for 5 minutes.
5. Place on magnetic stand and wait until liquid is clear (~5 minutes).
6. Remove and discard all supernatant from each well.
7. Wash beads as follows:
    a. Keep on magnetic stand and add 175 µL fresh 80% ethanol to each well.
    b. Wait 30 seconds.
    c. Remove and discard all supernatant from each well.
    d. NOTE: This protocol only uses a **single** ethanol wash.
8. Remove all residual ethanol from tube/well.
9. Air-dry on magnetic stand for 1 minute.
10. Remove from magnetic stand.
11. Add 10.5 µL of ELB (Elution Buffer).
12. Slowly pipette to mix until beads are fully resuspended.
13. Incubate at room temperature for 2 minutes.
14. Seal and centrifuge tubes/plate at 280 x g for 10 seconds.
15. Place on magnetic stand and wait until liquid is clear (~2 minutes).
16. Transfer 8.5 µL supernatant to a new plate/tube.
17. Add 8.5 µL EPH (Elute, Prime, Fragment High Mix) to each well.
18. Pipette up and down 10 times to mix.
19. Place on thermal cycle (run program above) and run.
    Each tube/well contains 17 µL of sample.

From this point, continue with the standard TruSeq Stranded Total RNA Human/Mouse/Rat library preparation protocol, starting at the 'Synthesize First Strand cDNA' step.

### Supplemental Figures

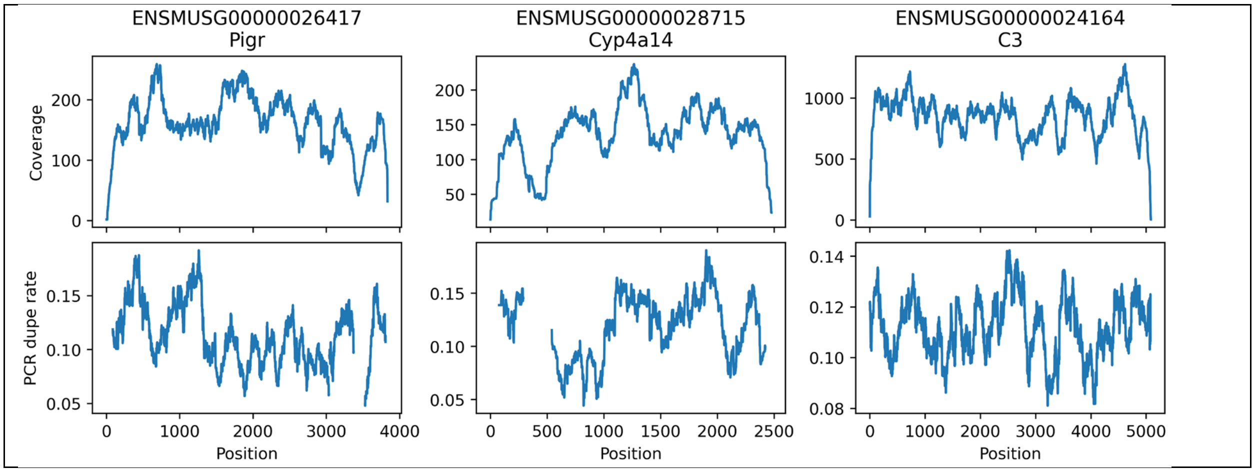


**<u>Figure S 1 - PCR dupe rate by position in NoSelect data</u>**

In NoSelect data from data set NS1, coverage depth (top row) and PCR dupe rate (bottom row). Overall PCR dupe rate is much lower in these experiments than in the PolyA experiments and the relationship between dupe rate and coverage is less clear. PCR dupe rate is shown only in regions where the coverage was at least 100, to avoid large statistical noise at low coverage.

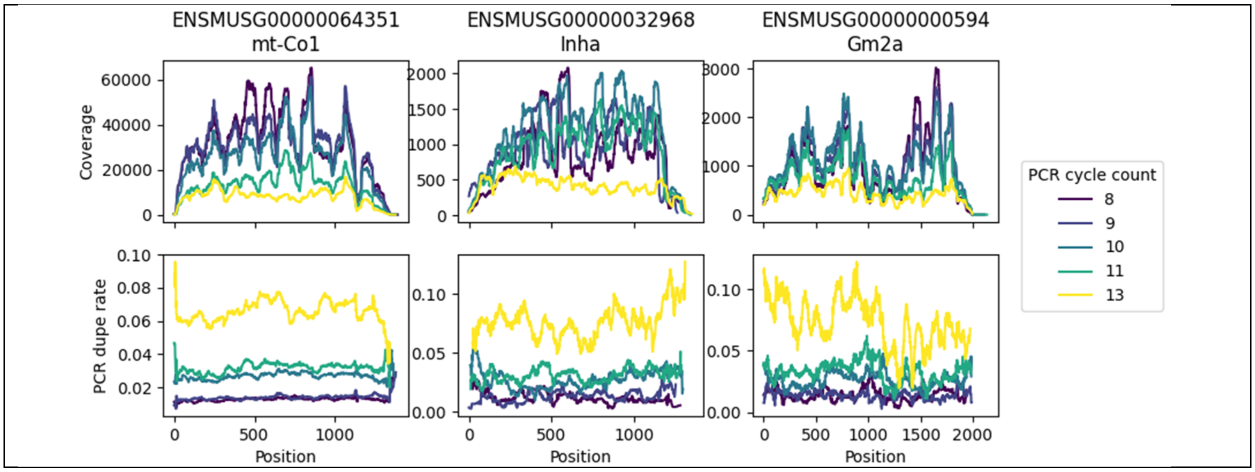


**<u>Figure S 2 - PCR cycle counts – absolute values</u>**

Mouse testis data from SRA accession PRJNA416930 with varying PCR cycle counts. Absolute coverage and PCR dupe rate are plotted by position. PCR dupe rate is determined by UMI tag and is computed as the fraction of deduplicated reads that originally had at least one duplicate. Samples with higher PCR cycle count also had lower input material.

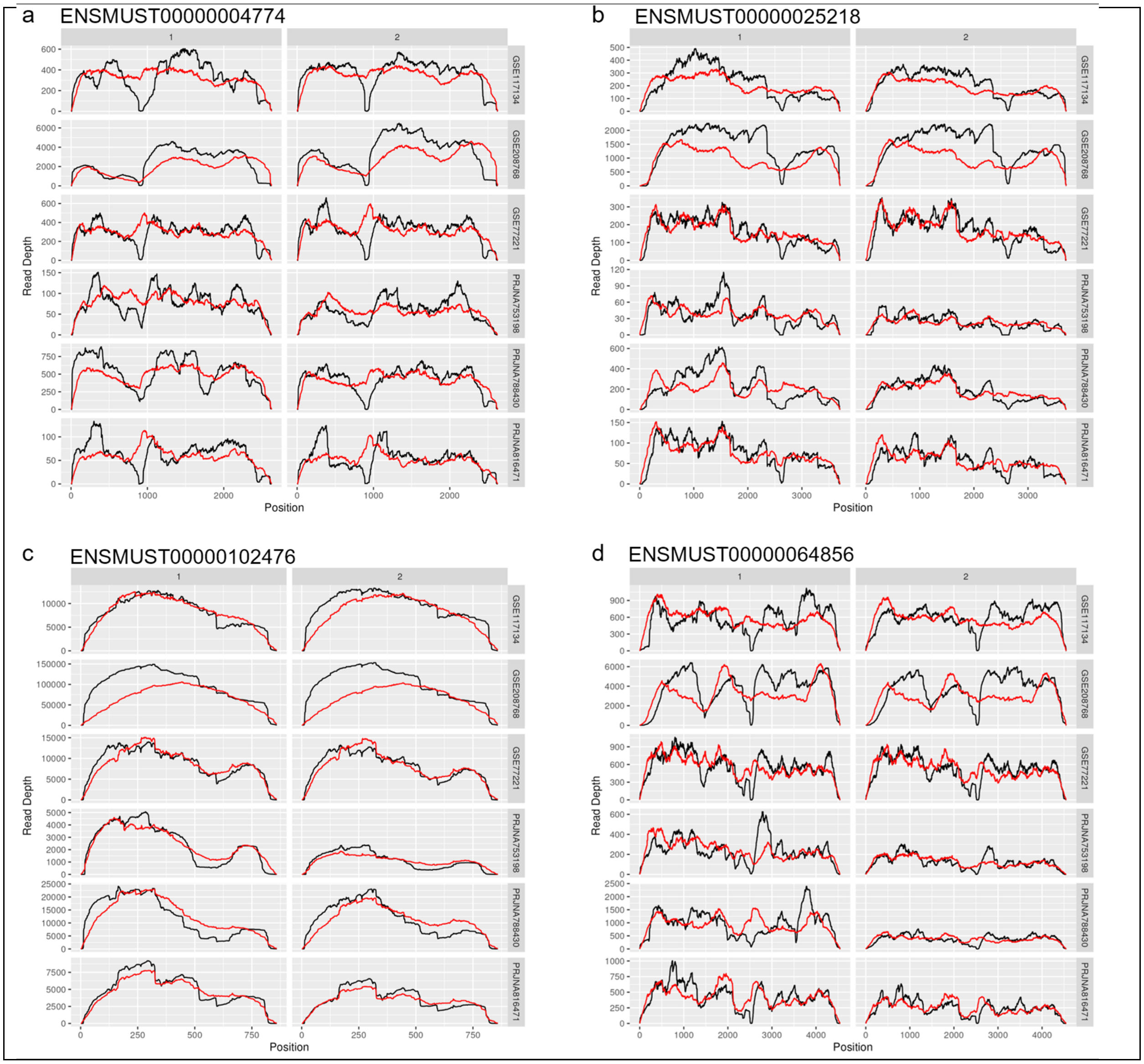


### Figure S 3 - Alpine fits

Alpine, a model of coverage non-uniformity, shown in red versus true coverage in black for four genes in two replicates (columns) and 6 studies (rows). Data set GEO used.

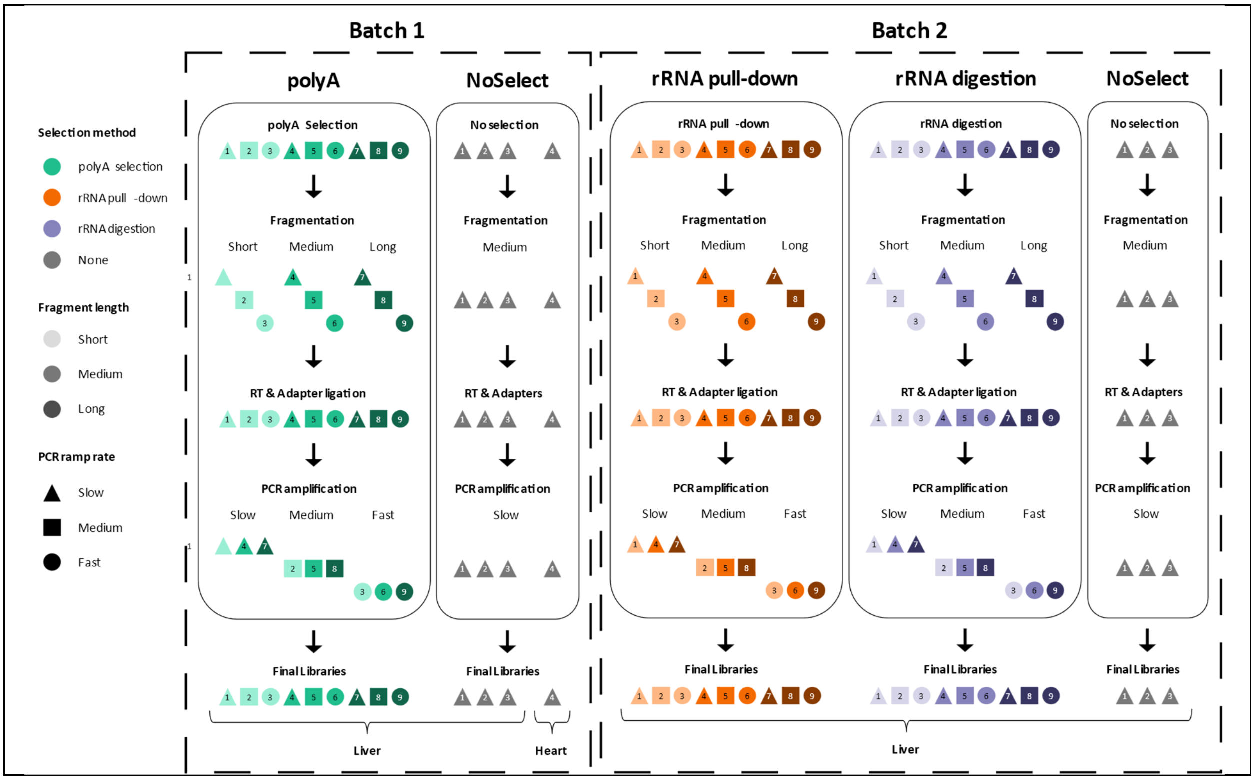


### Figure S 4 - Data set NS1 study design

For data set NS1, one mouse liver sample was sequenced in 27 libraries in two batches with all combinations of selection method (PolyA selection, rRNA pull-down, rRNA digestion), fragmentation length (short, medium, long) and PCR amplification ramp rate (slow, medium, fast). In addition, three liver libraries in each batch and one heart sample library were sequenced with the No Selection method, medium fragmentation length, and slow PCR ramp rate.
